# The role of migration in mutant evolution in fragmented populations

**DOI:** 10.1101/2021.06.09.447669

**Authors:** Jesse Kreger, Donovan Brown, Natalia L. Komarova, Dominik Wodarz, Justin Pritchard

## Abstract

Mutant evolution in fragmented populations has been studied extensively in evolutionary biology. With an increased focus on evolutionary dynamics in medical research, quantification of mutant load in fragmented populations with varying levels of migration has become especially important. Examples of fragmented populations are hematopoietic stem cell niches in the bone marrow where cells can re-circulate between niches through the blood, or colonic crypts where movement of cells across different crypts is not thought to be common. Here we use a combination of experiments and theory to investigate the role of migration in mutant distribution. In the case of neutral mutants, the experiments confirmed that while the mean number of mutants is not influenced by migration, the probability distribution is, which manifested itself in a change in the skewedness of the distribution of the mutant numbers in the demes. In the case of disadvantageous mutants, we investigated the phenomenon of the increase in the expected number of mutants compared to that of the selection-mutation balance. In a single deme, this increase is observed when the deme size is lower than the critical size, *N*_*c*_. In a fragmented system that consists of connected demes with a probability of migration, the increase in mutant numbers above the selection-mutation balance can be maintained in small (*N < N*_*c*_) demes as long as the migration rate is sufficiently small. The migration rate above which the mutants approach the selection-mutation balance decays exponentially with *N/N*_*c*_. These findings are relevant in the context of the complex and poorly understood processes that may lead to changes in the clonal composition in tissues and tumors.

## 1 Introduction

Understanding the principles of mutant evolution has been a major focus in evolutionary biology, and mathematical models have been an important component in this research. Different evolutionary measures have been considered, including the average number of mutants at a given time or population size, the fixation probability of mutants, and the average time to fixation for mutants of varying relative fitness [21, 43, 33, 18, 57]. Different types of evolutionary models have been explored, including the Moran process [38, 39] and the Wright-Fisher model [61] that assume constant population sizes.

The population structure is an important determinant of the evolutionary trajectories, and the dynamics on graphs [3, 2, 52, 4, 31], in spatially structured populations [1, 30, 40, 7, 32, 60], and in fragmented populations [16, 42, 54, 14] have been subject to investigation. Evolutionary dynamics in fragmented populations are of interest for questions connected to ecology and ecological conservation [53, 14, 28, 25, 45, 47, 17, 44, 51, 48, 8], but also have high relevance for the dynamics of cells in a biomedical context. Carcinogenesis is essentially an evolutionary process, and the emergence of various kinds of mutants has important clinical consequences, such as resistance to therapies. While solid tumors are often made up of a growing mass of cells, leukemias (such as chronic lymphocytic leukemia or CLL) can show more intricate population structures, where tumor cells grow separately from each other in different compartments of the hematopoietic system, including lymph nodes, the spleen, and the bone marrow. Although cell growth occurs in these distinct compartments, cells can also redistribute between them via the blood. This essentially corresponds to a fragmented population with migration. Cellular evolution in healthy tissues [27] can also occur in the context of fragmented populations. For example, the bone marrow, where hematopoietic stem cells reside, represents a complex environment where cells do not mix well but exist in niches that are spatially separated from each other. Hematopoetic stem cells home to a particular niche, and yet they also circulate systemically [56]. This process of homing and circulation resembles patch migration in fragmented populations. Other tissues can have similar characteristics, for example the colorectal tissue exists as a large collection of individual crypts, each of which contains approximately 5-10 stem cells [12], although cell migration between crypts is probably not a common process in this setting. The principles of somatic evolutionary processes in healthy and cancerous tissue are central to understanding how disease develops and how treatment resistance emerges.

Evolution in fragmented populations can be described by so called metapopulation models, which can also be referred to as deme, island, or patch models [39, 29, 14, 15, 28, 7, 48]. These models describe a group of distinct, spatially separated populations of the same type. Some amount of interaction between the separate groups occurs via migration of individuals from one group to another, and the dynamics within a single group of individuals is generally assumed to be nonspatial [16, 54, 29, 39]. Migration of individuals can either occur to the nearest neighboring regions (spatially restricted), or individuals can migrate to any region in the system. A higher rate of migration decreases population fragmentation because it results in each region’s dynamics becoming dependent on a larger portion of the overall population and thus in better mixing of individuals [15, 61, 39, 54].

Mathematical metapopulation models with different assumptions on structure and migration between groups have been studied in the context of evolutionary dynamics [16, 36, 6, 46, 54, 57, 11, 18, 50, 16]. Commonly, it is found that the fixation probability of a mutant is largely independent of migration (depending on the explicit model assumptions) [36, 37, 54], but that other quantities such as the time to fixation can vary based on model structure [50]. Processes related to extinction and recolonization of regions have been an area of interest in population ecology, see for example [14, 49, 53, 28, 48]. Furthermore, the field of adaptive dynamics has also been instrumental to understanding the evolution of metapopulations and effects of different migration strategies on population growth and persistence (see for example [42, 13]).

The interplay between metapopulation dynamics and traits or alleles has been previously studied in multiple experimental contexts [26, 20, 10]. For instance, Kerr et al identified path dependent migration effects in the eco-evolutionary dynamics of E.coli-T4 phage co-cultures [20]. Excitingly, recent work on range expansions in asexually reproducing microbes has show that an excess of spontaneous mutations (relative to Luria-Delbruck expectations) are generated during spatial range expansions by allele surfing [10]. However, to the best of our knowledge, no prior work has directly tested the distributions of existing neutral mutations across a large number of fragmented asexual populations in the presence versus absence of migration.

In this paper, we address two aspects of evolution in deme population structures that apply to somatic evolutionary processes: (i) What is the expected distribution of mutants across demes? This question, which we study both experimentally and theoretically, has relevance for assessing the clonal composition of such tumors in patients. For example, if neutral or disadvantageous mutants are likely to be present, together with wild-type cells, in most regions of the system, then sampling any of the regions will provide an accurate picture. An example is the screening for drug resistant mutants before the start of therapy. On the other hand, if some of the regions contain almost only mutant cells, and some of the regions contain almost only wild-type cells, then more complex sampling procedures will have to be employed to obtain an accurate picture of the genetic heterogeneity across the cell population. This has been demonstrated with spatially explicit computational models[62, 41], and is also relevant to fragmented, deme-structured populations. (ii) What is the effect of deme population structure on the selection-mutation balance of disadvantageous mutants, for example drug resistant mutants before treatment? The expected frequency of disadvantageous mutants at selection-mutation balance is well understood for homogeneous populations, but less so for deme populations structures. If the tumor cell population is fragmented into a collection of demes, does this benefit or hurt the population of disadvantageous mutants?

The paper starts with a description of our experimental findings, and these are interpreted with the help of a metapopulation model. We then go beyond the experimental setup and use the model to explore the evolutionary dynamics in the presence of de novo mutation processes and mutants of different fitness. Finally, we discuss how our insights can improve understanding of somatic evolutionary processes, especially in relation to carcinogenesis and cancer therapy.

## 2 Methods

### 2.1 Mathematical modeling

To study the role of population fragmentation and migration in evolution, we will consider a population of asexually reproducing individuals of two types, which we refer to as “wild types” and “mutants”, see Figure 1. The total population of *NK* individuals is split into *K* demes of *N* individuals each, as in for instance [16]. Within each deme, we model the stochastic birth-death dynamics by using the well-known Moran process (see e.g. [38, 39]). Therefore, the population of each deme as well as the total population remain constant. Competition is implemented by assuming that mutants may have fitness (denoted by *r*) that is not necessarily equal to the fitness of the wild types (assumed to be 1). Mutations are included through forward mutation (with probability *u* per division of a wild type cell) and back-mutation (with probability *u*_*b*_ per division of a mutant cell). Migration is modeled in the following way. A single migration event is attempted with probability 0 *≤ p*_*migr*_ *≤* 1 and performed by randomly selecting two demes, then randomly selecting *n*_*cells*_ cells from each and swapping them with each other. Prior to the Moran (birth-death) update for all the *K* demes, migration updates are completed *n*_*swaps*_ times. Table 1 lists all the model parameters; further details of the model are included in Supplementary Information Section 1.

**Figure 1:**
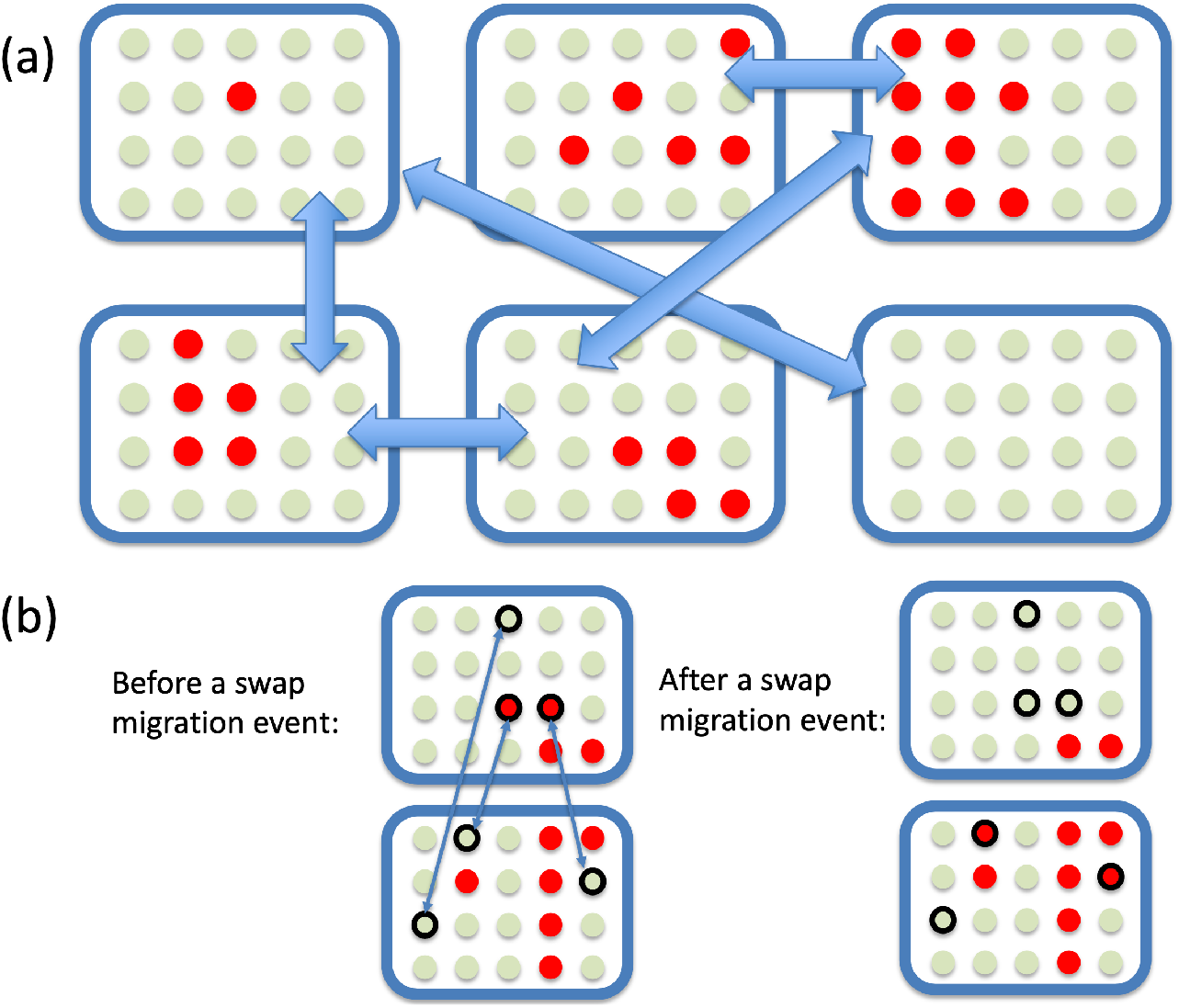
A schematic illustrating the mathematical model. (a) General model structure: each rectangle represents a deme, and green (red) circles represent wild-type (mutant) cells. Two-sided arrows represent random swap-migration events within randomly chosen pairs of demes. (b) Details of a swap migration event: groups of *n*_*cells*_ = 3 cells are randomly selected within two demes, and exchanged. As a result of this particular event, the number of mutants in the top deme decreased, and the number of mutants in the bottom demes increased by 2.

**Table 1:** Description of model parameters.

| Notation | Description |
| --- | --- |
| $K$ | number of demes |
| $N$ | constant number of cells in each deme |
| $NK$ | constant total number of cells in overall population |
| $r$ | fitness of the mutant cells |
| $u$ | rate of forward mutation wild type → mutant |
| $u_b$ | rate of back-mutation wild type ← mutant |
| $p_{migr}$ | migration probability |
| $n_{cells}$ | number of cells exchanged during a swapping event |
| $n_{swaps}$ | number of swaps that occur during a migration event |
| $j_{sel-mut}$ | selection mutation balance in each deme |

In the absence of forward mutation (*u* = 0), the model has only one absorbing state, given by mutant extinction in all demes. Similarly, in the absence of back mutation (*u*_*b*_ = 0), mutant fixation in all demes is the only absorbing state. With the inclusion of both forward and back mutation, there are no absorbing states [38]. In the absence of migration, the stationary probability distribution in an individual deme can be calculated. For example, in the regime where mutant fixation happens on a much faster time-scale than mutant production, a simple expression for the stationary probability can be derived. Denoting by *y*_*i*_ the probability to have *i* mutants in the deme, we have

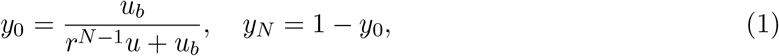

with the probability of the other states being of the order of the mutation rate. This is similar to previous approximations of the stationary distribution for the Moran process with mutation and selection and under various conditions, see for instance [39, 9, 55, 19, 54]. Another useful quantity is the selection-mutation balance, which is given (for an individual deme, in the limit of small mutation rates) by

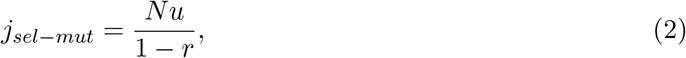

this quantity represents the number of (negatively selected, *r <* 1) mutants, in the Moran process with mutations, which corresponds to an equal probability to increase and decrease this number in a single birth-death update. Supplementary Information Section 2 (see also [9, 55, 39]) provides details of the calculations for expressions (1) and (2), as well as higher order approximations.

### 2.2 Experiments with neutral mutants

To address the existing gap in the literature concerning studies of neutral mutant dynamics in fragmented populations in the presence and absence of migration, we performed experiments that represent an *in vitro* comparison to the mathematical model presented in the preceding section. To create a system representing neutral migration we mixed GFP labeled mammalian cells with unlabeled cells. This suspension of mammalian cells could be propagated in the wells of a 96 well plate. By systematically transferring small volumes of the cell suspension, we experimentally simulated migration between wells. Cells were continually maintained at confluent cell population densities to mimic the Moran process. The proportion of mutants in the wells both with and without migration of cells between wells was assessed at the end of the experiment. Further details on the *in vitro* experiments are presented in Supplementary Information Section 3.

## 3 Results

### 3.1 Population fragmentation changes mutant distribution for neutral mutants

To motivate this study, we performed some simple experiments that examined the role of fragmentation and migration in neutral mutant dynamics. 96 wells were filled with cells, such that each well contained 1% of (neutral) mutants. The process of cell migration was implemented by swapping a small percentage of cells between demes by using a pipette. The number of mutants in the wells was assessed at the end of the experiment and compared with the control condition with no migration. The resulting experimentally obtained distribution of the mutant numbers is shown in Figure 2. We observed that while the mean number of mutants in the absence and in the presence of migration was the same, the distribution was significantly different; in particular, the distribution without migration had a much larger skewness, while in the presence of migration it was more symmetric.

**Figure 2:**
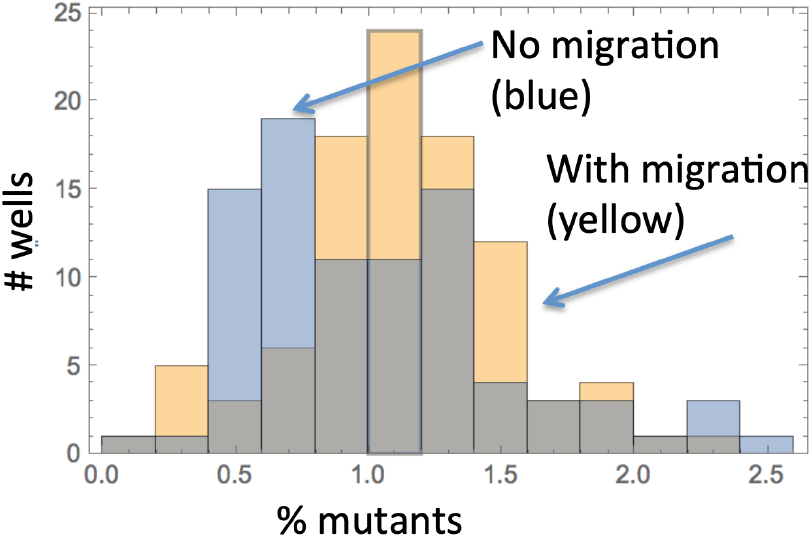
Effect of migration on neutral mutant distribution, experimental results. The blue bars represent the control condition without migration between the wells and the yellow bars represent the experimental condition with migration. Initial condition of 1% mutants in each well. The average percent of mutants in each well without migration is 1.03%, and with migration is 1.13% (not significantly different using T-test, *p*-value greater than 0.1). The Kolmogorov-Smirnov test between the two distributions gives a *p*-value of about 10^*−*3^, which suggests that the distributions are significantly different. The skewness without migration is 0.89, and with migration it is much smaller at 0.07.

To explain these observations and extend the results to other conditions, we began by analyzing the dynamics of neutral mutants (*r* = 1).

#### Unimodal versus bimodal mutant distribution in the absence of mutations

To reproduce the experimental set-up, we assumed that there is no mutation, and started with some small initial number of mutants in each deme (*m*_0_). In order to analyze the effect of migration/population fragmentation, we ran simulations with and without migration of cells between the distinct demes.

We found that the mean percent of neutral mutants is independent of migration, and is equal to the initial ratio of mutants in the system. This is because the transition probabilities are symmetric, see Supplementary Information Equation (1) and Section 4.1 for details. Furthermore, as the wild type and mutant are neutral, the probability for the mutant to fixate within the system is equal to the initial frequency of mutants in the system (although the time to such fixation depends on migration [50, 57]). This is because in the context of a symmetric random walk, the fixation (i.e. absorption at the upper boundary) probability is proportional to the initial condition, see Supplementary Information Section 4 and [36, 38, 39].

On the other hand, the distribution of the number of mutants in the system and the dynamics within individual demes are significantly influenced by the presence of migration (Figure 3). Starting from a delta-like distribution (as initially all demes contain *m*_0_ mutants), the distributions get wider with time and eventually reach a quasi-stationary distribution. In the absence of migration (panel (a)), the dynamics of each deme are independent from one another. Since the probability of fixation is simply the initial fraction of mutants (*m*_0_*/N*), there is a large chance (given by 1 −*m*_0_*/N*) of mutant extinction in each deme. Therefore, the probability distribution of the number of mutants in each deme becomes flatter and develops a skew to the right, as most demes will trend toward mutant extinction, while a few will trend toward mutant fixation. In the presence of migration (panel (b)), the dynamics of each deme are no longer independent from one another. As was similarly found in [54, 16, 15, 39, 61], migration makes all the demes look more homogeneous to each other, resulting in a one-humped (unimodal) distribution. These results match well with *in vitro* experimental simulations of the computational model, which are shown in Figure 2. Note that while the figure shows a long-term state, this is not an equilibrium, and the only outcomes as time → ∞ is mutant fixation or mutant extinction in the whole system [16, 39].

**Figure 3:**
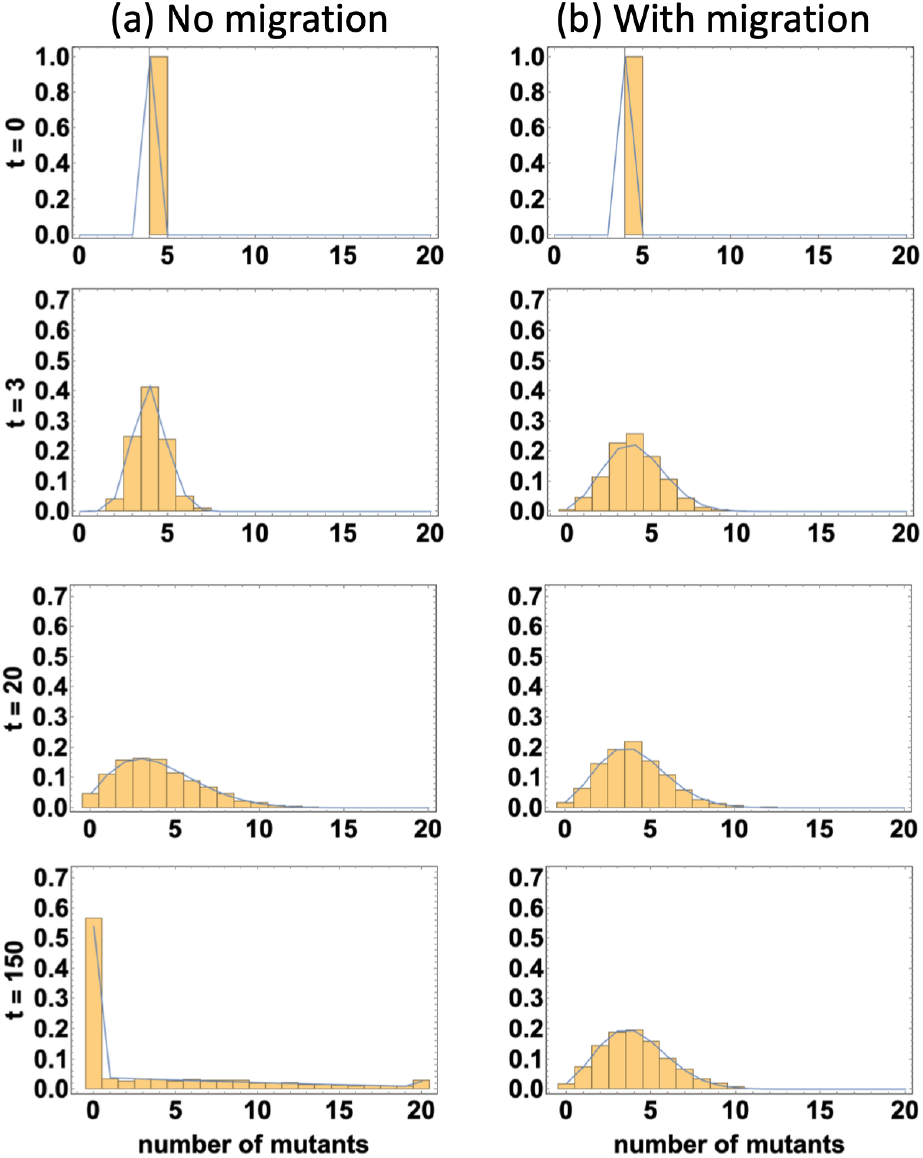
Stochastic simulations (histograms) and iterations of Equation (2) in the Supplementary Information (blue lines). Panel (a) represents the absence (*p*_*migr*_ = 0) and panel (b) represents the presence (*p*_*migr*_ = 1) of migration. The probability distributions are presented at several moments of time (*t* in each plot corresponds to the number of discrete Moran steps). The rest of the parameters are *N* = 20, *m*_0_ = 4, *n*_*swaps*_ = 750, *n*_*cells*_ = 5, *r* = 1, *u* = *u*_*b*_ = 0, and *K* = 1.5 *×* 10^3^.

In addition to the extremes of no migration or a large amount of migration (where the system is well-mixed), we also investigate other regimes where there is some intermediate level of migration of cells between the demes in the system. Figure S3 shows the time-evolution of the mutant probability distributions obtained by iterating Equation (2) in the Supplementary Information (see panels (a-c) for three different values of *p*_*migr*_), and then by plotting the resulting quasi-stationary probability distributions (panel (d)). Here we see that the effect of the Moran process is to “make” the probability distribution bimodal, and the effect of migration is to “make” it unimodal. The result is a trade-off of the two tendencies, and depending on the amount of migration, the distribution shape changes accordingly.

#### Quasi-stationary distributions become stationary in the presence of mutation

Next, we expand the theory beyond the experimental conditions of Figure 2 to include the effect of mutations. Since mutants are now generated stochastically, we alter the initial conditions to start with the wild type fixated in all demes (*m*_0_ = 0). In the case of only forward mutation, the mutant will be effectively advantageous and will fixate quickly in the entire population (see Figure S5(a-b)). On average, the time to fixation decreases with increasing migration, as faster migration results in more frequent introduction of the mutant into all of the demes, see [54, 50, 14, 57, 50].

In the case of both forward and back mutation, the dynamics are more complex. Figure 4(a) shows simulations representing 2 × 10^3^ demes of 20 cells each, after 10^5^ iterations with varying rates of migration. The histograms represent the number of neutral mutants per deme. In the absence of migration (left), the number of mutants will drift around, becoming extinct or fixated within a deme. In a highly fragmented population, this will happen more often, and the mutant will be at the extinction/fixation long-term state most of the time. Increasing migration (and/or decreasing population fragmentation by increasing the size of the demes, not shown) will result in fluctuation around the selection-mutation balance in each deme (Figure 4(a, right)). As in the simulations without mutation, if the rate of forward and back migration is equal (*u* = *u*_*b*_), then migration does not change the expected mean number of neutral mutants (as the stationary distribution is symmetric around the selection-mutation balance of 50% mutants, see Figures S4 and S5(c-d)). If the rate of forward and back migration is not equal (*u* ≠ *u*_*b*_), then the expected mean number of mutants and the selection-mutation balance is simply 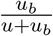 [39].

**Figure 4:**
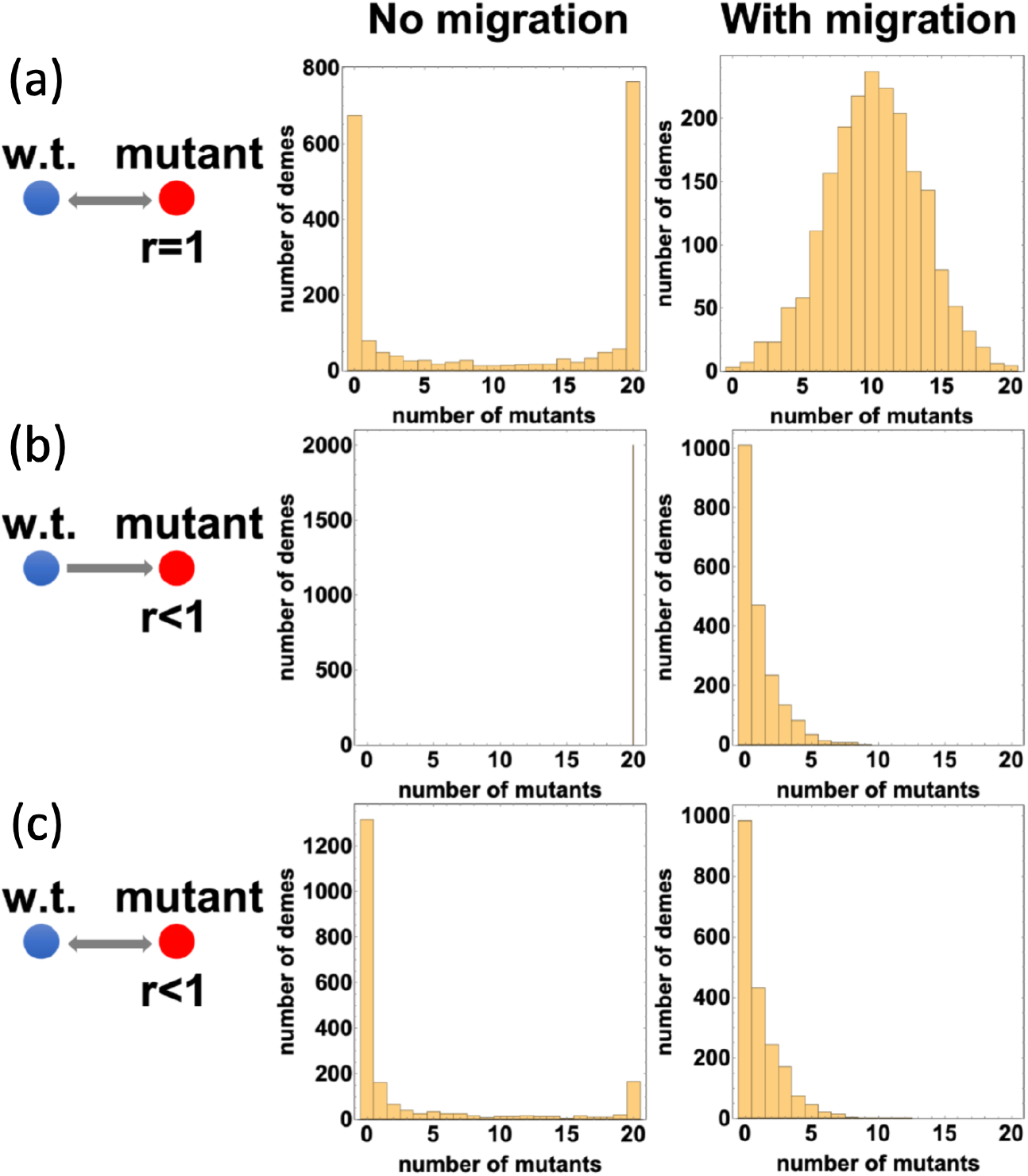
Histograms for the number of mutants per deme in the absence (*p*_*migr*_ = 0) and presence (*p*_*migr*_ = 1) of migration, after 10^5^ Moran iterations. (a) Neutral mutant (*r* = 1) with both forward and back mutation (*u* = *u*_*b*_ = 0.005), see Figure S4 for intermediate migration cases; (b) disadvantageous mutant (*r* = 0.9) with forward mutation only (*u* = 0.005, *u*_*b*_ = 0), see Figure S6 for intermediate migration cases; (c) disadvantageous mutant (*r* = 0.9), with forward and backward mutation (*u* = *u*_*b*_ = 0.005), see Figure S7 for intermediate migration cases. The horizontal axis is the number of mutants and the vertical axis is the number of demes at that number of mutants. Other parameters are *N* = 20, *m*_0_ = 0, *n*_*swaps*_ = 100, *n*_*cells*_ = 10, and *K* = 2 × 10^3^.

Note also that the presence of mutations changes the nature of the long-term system behavior: the quasi-stationary distribution observed in the absence of mutations (Figure 3(b), bottom graph) becomes a stationary distribution in the presence of forward and back mutation, as absorbing states no longer exist [16].

### 3.2 Population fragmentation changes mutant numbers and distribution for disadvantageous mutants

Next we turn to the dynamics of disadvantageous mutants. While this scenario is highly biologically realistic for cell populations, it is more difficult to study experimentally due to the lower probability of mutant growth. In the absence of mutation, a small initial number of disadvantageous mutants will likely decay quickly and go extinct. Therefore, we focus on mathematical models that include mutation processes. We will show that while migration changes the distribution of demes in a similar manner for both disadvantageous and neutral mutants, in the disadvantageous case migration also changes the expected number of mutants at the (quasi)-stationary state in fragmented populations. This is related to the concept of “drift load” [34, 35, 59] and “mutational meltdown” [58, 24], which describe how the accumulation of deleterious mutations can cause a gradual reduction in population size (and in small populations random genetic drift will progressively overpower selection making it easier to fix future mutations) potentially leading to population extinction. As we assume constant population sizes, which is relevant for healthy tissue dynamics or temporary steady states during tumor development, size fluctuation and extinction cannot occur; instead we observe elevated fractions of disadvantageous mutants depending on migration and population structure.

#### Fragmentation increases mutant numbers and decreases time to fixation

In the absence of back mutations, mutant fixation in all demes is the only absorbing/stationary state, which will again eventually be reached with 100% probability. However, when there is a large amount of migration and/or a large, well-mixed population, then fixation will take a very long time and quasi-stationary states are possible [16, 50].

Figure 4(b) shows a system of small patches in histogram form in the absence and presence of migration, for disadvantageous mutants with only forward mutation. When the overall population of cells is highly fragmented (no migration, left), fixation will occur quickly in each of the individual demes, and thus in the overall population as well. However, if the overall population of cells is well-mixed, then fluctuation around a quasi-stationary state that is equal to the selection-mutation balance in each deme is observed (panel (b, right)). Between these extreme scenarios, we observe that the overall system fluctuates around a quasi-stationary equilibrium that is between selection-mutation balance in each deme and complete fixation in the overall system (see Figure S6). We can see that in the case of disadvantageous mutants, population fragmentation does not only change the distribution of mutants, but also increases the expected number of disadvantageous mutants. To further illustrate this, Figure 5(a-b) shows the time course of the number of disadvantageous mutants for different rates of migration. Here we can see the (quasi)-stationary number of mutants in the system, and that there are on average more mutants expected with lower migration rates (higher levels of population fragmentation), because fixation in each deme is more easily reached for fragmented (small) populations [58, 35].

**Figure 5:**
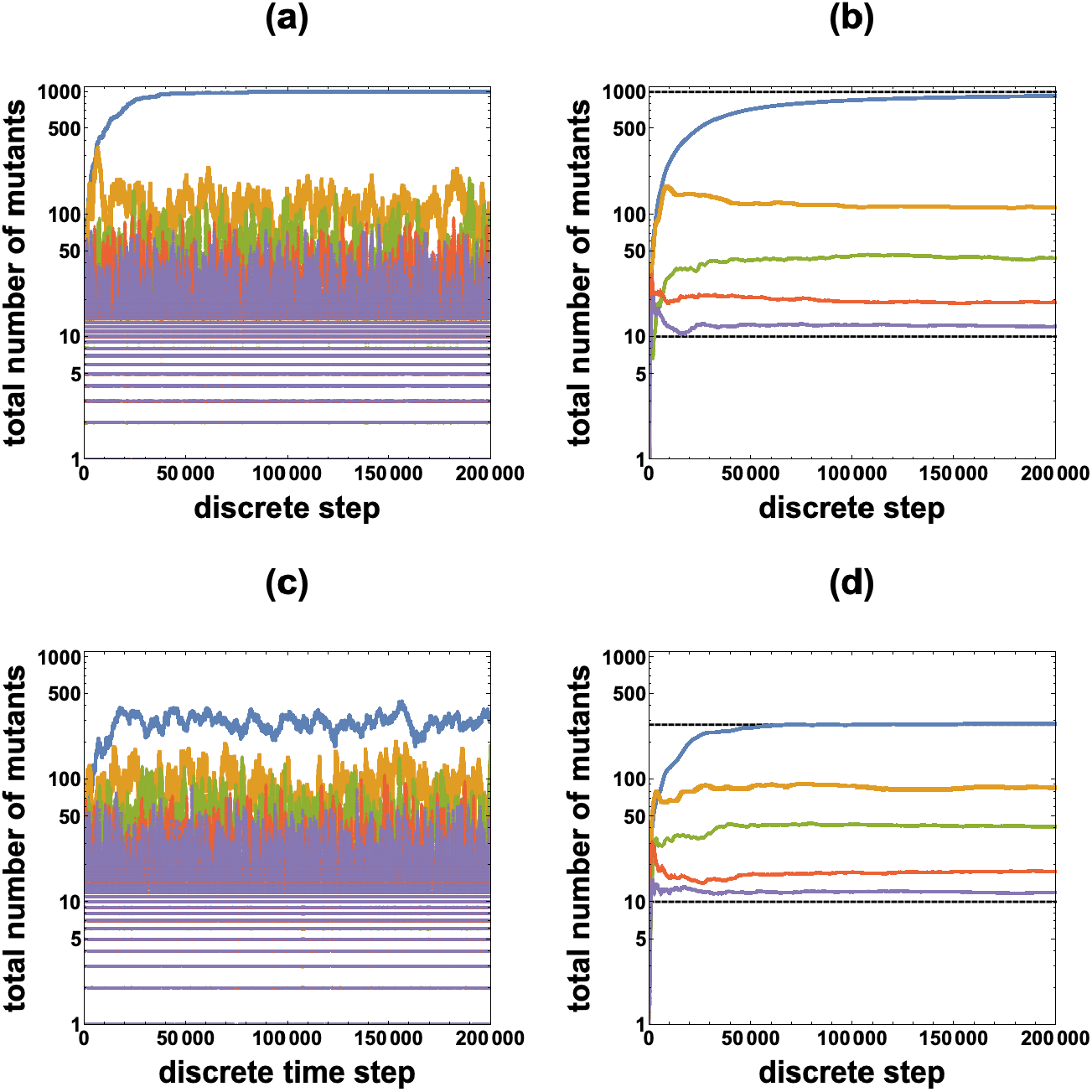
Number of mutants over time for varying rates of migration with a disadvantageous mutant (*r* = 0.9). Other parameters are *N* = 10, *m*_0_ = 0, and *K* = 100. Selection-mutation balance is approximately 10 mutants in the system and mutant fixation is 10 mutants in each deme. Blues lines, no migration. The approximate expected number of mutants can be calculated using Equation (1). Yellow lines, low migration: 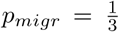, *n*_*swaps*_ = 1, and *n*_*cells*_ = 1. Green lines, medium migration: *p*_*migr*_ = 1, *n*_*swaps*_ = 1, and *n*_*cells*_ = 1. Red lines, high migration: *p*_*migr*_ = 1, *n*_*swaps*_ = 5, and *n*_*cells*_ = 1. Purple lines, very high migration: *p*_*migr*_ = 1, *n*_*swaps*_ = 10, and *n*_*cells*_ = 5. The approximate expected number of mutants is the selection-mutation balance. (a) Forward mutation only (*u* = 10^*−*3^, *u*_*b*_ = 0), number of mutants in the system at each time step. (b) Forward mutation only (*u* = 10^*−*3^, *u*_*b*_ = 0), temporal average of the number of mutants in the system at each time step. Dashed lines represent the selection-mutation balance and mutant fixation. (c) Forward and back mutation (*u* = *u*_*b*_ = 10^*−*3^), number of mutants in the system at each time step. (d) Forward and back mutation (*u* = *u*_*b*_ = 10^*−*3^), temporal average of the number of mutants in the system at each time step. Dashed lines represent the selection-mutation balance (Equation (2)) and the predicted expected number of mutants under no migration (Equation (1)).

The overall dynamics for both individual demes and for the total number of disadvantageous mutants with varying levels of migration and deme size and only forward mutation are summarized in Supplementary Table 1.

#### Fragmentation increases mutant numbers even when fixation is not an absorbing state

In the case of both forward and back mutations, there are no longer any absorbing states.

Figure 4(c) shows histograms for the number of disadvantageous mutants in the absence and presence of migration, with the inclusion of back mutation. The dynamics are similar to the forward mutation only case (panel (b)), except the quasi-stationary distributions described in the preceding paragraph are now stationary distributions, as demes will not all eventually trend toward fixation (left panel). In particular, depending on the level of population fragmentation, demes will either fluctuate around a stationary value, or will individually bounce back and forth between mutant extinction and mutant fixation. In the latter case (high fragmentation), the system is characterized by a higher expected number of mutants compared to the well-mixed (or high migration rate) system. As seen in Figure 5(c-d), since mutant fixation is no longer an absorbing state, we expect a smaller number of mutants compared to when there is only forward mutation (panels (a-b)). The expected number of mutants in the absence of migration can be computed in the case of a small mutation rate, according to Equation (1). As the amount of migration increases, the expected number of mutants converges to the selection-mutation balance given by Equation (2).

The overall dynamics for both individual demes and for the total number of disadvantageous mutants with varying levels of migration and deme size and both forward and back mutation are summarized in Figure 6 (see also Supplementary Information Table 2).

**Figure 6:**
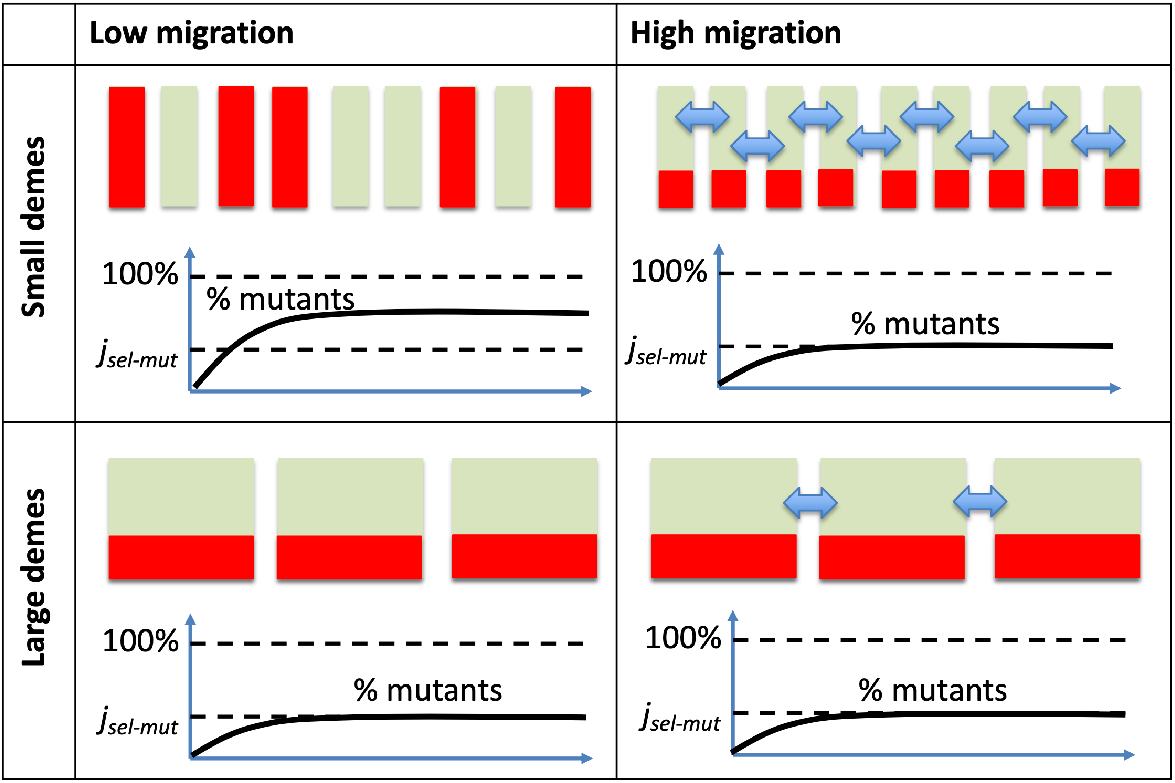
Summary of results: forward and back mutation with varying migration (columns) and deme size (rows), assuming a constant total population. There are no absorbing states. Individual demes are represented as green rectangles, and the level of mutants in each is shown in red. Total number of mutants panels schematically show the percent of mutants as a function of time; the black dashed lines represent the selection-mutation balance, *j*_*sel*−*mut*_, and 100% fixation. For a text description of this figure, see Supplementary Table 2.

#### When can we expect to see more mutants than predicted by selection-mutation balance?

The amplification of the number of disadvantageous mutants in fragmented populations requires the population to be highly fragmented (that is, the individual patches must be sufficiently small) and the migration rate to be not too high, see figure 6.

Even in the absence of migration, if each deme size is too large, then fixation will almost never be reached and fluctuation around the selection-mutation balance in each deme will be observed instead. On the other hand, if the deme size is very small (*N* = 2), then the expected number of mutants is approximately 50% of the system, as each deme will spend about 50% of the time at mutant extinction and 50% of the time at mutant fixation because of the small mutation rate. As the number of cells per deme increases, this effect of fixation continues to increase the number of mutants, but contributes less and less as the fixation probability decreases. Therefore, as the deme size increases, the expected number of mutants (the red line in figure 7) will extrapolate between two regimes: (i) the fast fixation regime, where the mean number of mutants in a deme given by *Ny*_*N*_ (equation (1), green line in figure 2), and (ii) the selection-mutation balance (equation (2), yellow line in figure 7). To estimate the threshold deme size, *N*_*c*_, above which the expected number of mutants becomes close to selection-mutation balance, we find the intersection of the fast-fixation (green) and selection-mutation balance (yellow) lines by solving the equation *Ny*_*N*_ = *j*_*sel−mut*_ for *N* :

**Figure 7:**
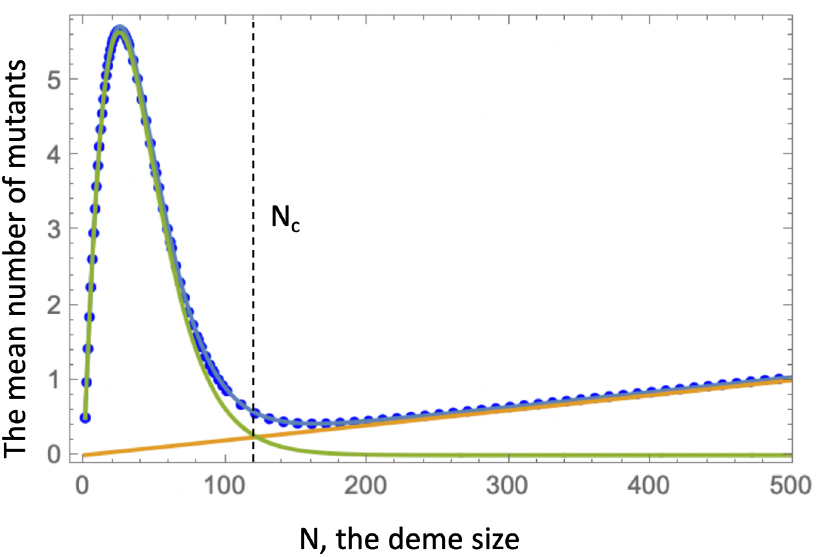
Estimating *N*_*c*_. The expected number of mutants in a single deme in the absence of migration is shown as a function of *N* ; it is computed numerically (blue circles) by determining the principal eigenvector of the transition matrix, see Equation (1) of the Supplementary materials, and also by using approximation (1) and (8), Supplementary materials (blue line). The green line represents the fast fixation regime (*Ny*_*N*_, equation ((1)); the orange line is the selection-mutation balance, *j*_*sel*−*mut*_ (equation (2)). The parameters are *u* = *u*_*b*_ = 10^*−*4^, *r* = 0.95. The threshold value *N*_*c*_ is shown by the dashed vertical line.

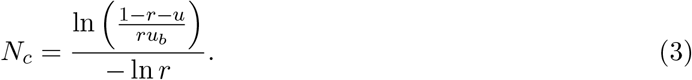

If the deme size is smaller than *N*_*c*_, a significantly larger number of mutants compared to the selection-mutation balance is expected. If however demes are connected to each other and migration is present, this may result in a lowering of the overall number of mutants. Under intense migration, the expected number of mutants tends to that predicted by selection-mutation balance. Therefore, an important question is the level of migration that is sufficient to lower the mutant levels back to that of election-mutation balance.

In this model, the overall intensity of migration (monotonically) depends on several parameters (see table 1): the probability of a migration event per update (*p*_*migr*_), the number of swaps during a migration event (*n*_*swaps*_), and the number of cells exchanged during a swapping event (*n*_*cells*_). To simplify the discussion, we will fix two of these to *n*_*swaps*_ = *K/*5 and *n*_*cells*_ = *N/*5, focusing on the parameter *p*_*migr*_ as the one parameter determining the rate of migration.

Figure 8(a) demonstrates how a threshold value of the migration probability can be calculated. Fixing the values of *u, u*_*b*_, and *r*, simulations were run for different choices of the deme size, *N < N*_*c*_, and the mean concentration of mutants (that is, the mutant number divided by the total population size, *NK*) was determined for each *p*_*migr*_. As anticipated, the expected mutant numbers are higher than the level predicted by the selection-mutation balance; also, they decrease with the deme size, *N*, and migration probability, *p*_*migr*_. To quantify the migration probability that, for each *N*, corresponds to a significant decay in the mutant population, we defined *p*_*c*_ as the value of *p*_*migr*_ that leads the number of mutants to fall to twice the selection-mutation balance. In figure 8(a), intersections of the mutant numbers with 2*j*_*sel−mut*_ are marked with colored symbols and their horizontal coordinate gives *p*_*c*_. This quantity decreases with *N*.

**Figure 8:**
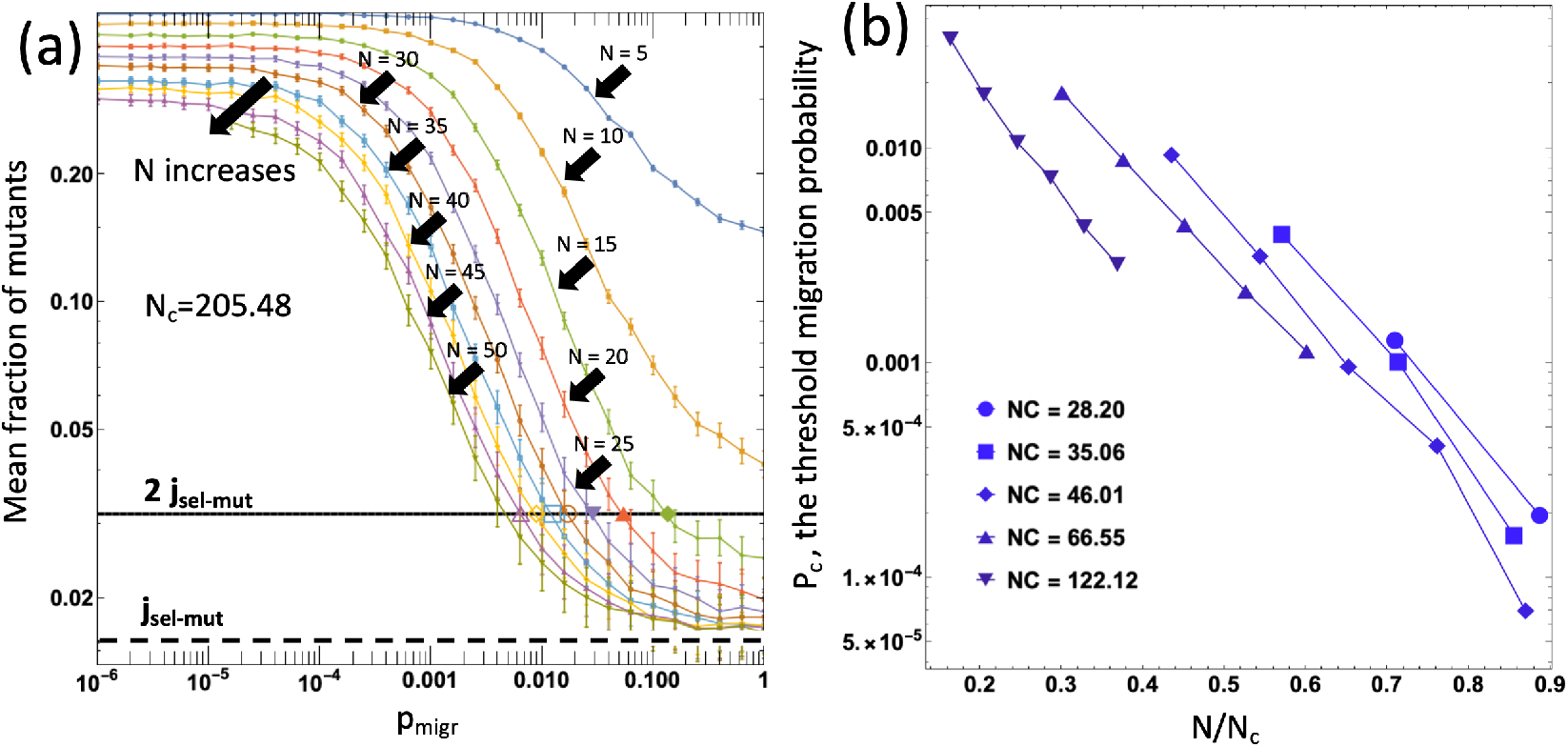
The role of migration in the level of mutants. (a) The mean number of mutants as a function of *p*_*migr*_, calculated as a temporal average over 10^8.5^ time-steps; the bars represent the standard error. Different curves correspond to different values of *N*. The horizontal dashed and solid lines are *j*_*sel−mut*_ and 2*j*_*sel−mut*_, respectively. The parameters are *u* = *u*_*b*_ = 10^*−*3.5^, *r* = 0.98, with *N*_*c*_ = 205.48. (b) The threshold values, *p*_*c*_, are plotted against the corresponding *N/N*_*c*_, for several values of *N*_*c*_. The exponent *B* (equation (4)) is 10.7 *±* 0.9. The rest of the parameters are: *K* = 20, *n*_*swaps*_ = *K/*5, *n*_*cells*_ = *N/*5.

Figure 8(b) shows the threshold migration rate as a function of *N/N*_*c*_ for several different values of *N*_*c*_. We observe that the dependence is exponential, and propose the empirical law:

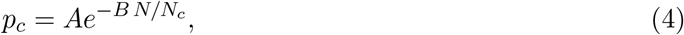

where the constants *A* and *B* do not depend on *N*.

## 4 Discussion and Conclusion

We have investigated the effect of migration on the mutant distribution in the demes of a fragmented system. In the case of neutral mutants, we performed experiments with cell colonies grown in 96 wells, where migration was implemented as swapping small numbers of cells between randomly chosen wells with a pipette. The experimental findings confirmed that while the mean number of mutants is not influenced by migration, the probability distribution is, consistent with theoretical predictions.

In the case of disadvantageous mutants, we investigated the phenomenon of the increase in the expected number of mutants compared to that of the selection-mutation balance. In a single deme, this increase is observed when the deme size is lower than the critical size, *N*_*c*_, given by equation (3). In a fragmented system that consists of connected demes with a probability of migration, the increase in mutant numbers above the selection-mutation balance can be maintained in small (*N < N*_*c*_) demes as long as the migration rate is sufficiently small. The migration rate above which the mutants approach the selection-mutation balance decays exponentially with *N/N*_*c*_, see equation (4).

These results demonstrate that deme structure and details about the migration processes can have important consequences for patterns of mutant evolution, which in turn can be of practical relevance. The dynamics explored here can be applied to two types of settings. First, consider tumors where cells grow in a deme-structured manner. Although tumors grow over time and we have considerd the evolutionary dynamics in constant populations, tumors can be characterized by periods of slow growth or temporary stasis until further mutants are generated that allow the cells to overcome specific selective barriers. Examples might be early colorectal adenomas, where cells are organized as glands that resemble crypt structures in the corresponding healthy tissue, or slowly growing / indolent cases of chronic lymphocytic leukemia, where cells grow in spatially separated lymph nodes, the spleen, and the bone marrow. The evolution of drug-resistant mutants is a major problem that results in the eventual failure of therapies, and the level at which resistant mutants pre-exist before the start of treatment tends to be an important determinant of the time to disease relapse [22, 5]. Mathematical models have been used to calculate the number of drug-resistant mutants, for example in chronic lymphocytic leukemia [22] or chronic myeloid leukemia [23], and these models assumed a spatially homogeneously growing tumor cell population. If drug-resistant mutants carry a fitness cost and are therefore disadvantageous before the start of therapy, the models analyzed here indicate that the organization of cells into demes can have a significant influence on the level at which such mutants pre-exist. Depending on the deme size the and migration rate, the number of resistant cells can be significantly larger than predicted by the selection-mutation balance. Beyond resistant mutants, our analysis has shown that deme structure and the rate of cell migration can impact the distribution of mutants across the demes, which has implications for understanding the genetic composition of the tumor cell population as a whole.

Healthy tissues at homeostasis correspond more precisely to our model in which populations are constant. Clonal evolution takes place within tissues as individuals age, and this has been clearly documented in the hematopoietic system [27]. In the condition called clonal hematopoiesis of indeterminate potential, mutant cell clones emerge in healthy individuals that are typically associated with a malignancy. These evolutionary processes take place in the bone marrow, where stem cells exist in niches, with traffic between different parts of the bone marrow via the blood [56]. According to our model, this deme structure can influence the exact spatial genetic composition of the cell population, with mutants being dominant in some parts of the bone marrow but not others. In the model, the details depend on the local population size and the migration rate of cells between the demes. As mentioned above, colonic crypts also represent a situation in which stem cells are fragmented across many demes with relatively small cell population sizes. In contrast to the hematopoietic system, however, movement of cells from one crypt to another is probably not a frequent occurrence in this case, which would lead to significantly different patterns of mutant distribution across the demes. This analysis thus highlights parameters that are important to measure to better understand the evolutionary dynamics in different tissues.

Finally, we note that in our model analysis, migration is assumed to be between random demes (i.e. not spatially restricted). For migration that is spatially restricted, disadvantageous mutant levels will be elevated compared to non-spatially restricted migration because once a region of demes becomes fixated with the mutant, it is less likely that the wild type will be reintroduced due to spatial restrictions. Therefore, spatially restricted migration increases population fragmentation compared to non-spatially restricted migration, which increases the number of mutants. Including different spatially restricted patterns of migration could be an interesting extension of our current work.

